# Chronic social stress induces isolated deficits in reward anticipation on a neuroeconomic foraging task

**DOI:** 10.1101/2022.01.17.476514

**Authors:** Romain Durand-de Cuttoli, Freddyson J. Martínez-Rivera, Long Li, Angélica Minier-Toribio, Scott J. Russo, Eric J. Nestler, Brian M. Sweis

## Abstract

Measuring reward anticipation distinct from other aspects of reward value, including costs required to obtain a reward or the intrinsic hedonic value of consuming the reward itself, can be difficult to disentangle. Here, we show that mice trained on a self-paced neuroeconomic foraging task convey reward anticipation via differentially invigorated travel times between uniquely flavored feeding sites separate from willingness to wait, consummatory behaviors, or place preferences measured within the same trial. Following exposure to chronic social defeat stress, we found that only stress-susceptible but not stress-resilient mice revealed deficits in this metric after consuming but not after rejecting a reward on the previous trial, indicating that blunted anticipation in these animals is state-dependent, or punctuated by recent reward receipt. After increasing economic pressure and task demands, locomotion was globally invigorated and, in turn, masked stress-related deficits in reward anticipation. These findings suggest that the ability to detect changes in specific aspects of motivational deficits associated with depression and other stress-related disorders depends on an interaction between the state of an individual and environmental circumstances.

---

Anhedonia describes a loss of interest or pleasure in rewarding activities and can manifest in the form of decreased motivation to engage in reward-seeking behaviors (*1*). Deficits in reward anticipation comprise one aspect of anhedonia. Levels of motivation, however, are highly dynamic and can be altered, for example, due to fluctuations in different physiological states of an individual or as a result of environmental circumstances (*2*).

Animal studies operationalize anhedonia in various ways, including decreased sucrose preference, impaired conditioned place preference, diminished effort expenditures on operant-based tasks (e.g., lever pressing or nose poking for food), blunted sensitivity on probabilistic reward tasks, reduced willingness to wait on delayed discounting tasks, and even alterations in orofacial expressions following reward consumption (*2–11*). Such experimental approaches, while instrumental in laying a foundational understanding of the neurobiology of motivation, include aspects of reward anticipation that are inherently confounded by hedonic processes related to the appraisal of the consumption of the reward itself or are directly linked to variations in the cost required to obtain the reward. It is therefore difficult to disentangle multiple properties of reward value from isolated, independent measures of intrinsic motivation pertaining solely to the process of reward anticipation.

Here, we tested mice previously exposed to chronic social defeat stress, a well-validated animal model used for the study of depression (*12–15*), on a translational neuroeconomic self-paced foraging paradigm, termed “Restaurant Row” (*12–21*). This task provides a rich framework to operationalize reward value and motivation-related processes among several domains of complex behavior separated across space and time. By analyzing travel behavior between reward patches, we found that the speed with which mice traveled was invigorated when approaching more preferred rewards – a metric of reward anticipation that was independent of other aspects of reward value measured within the same trial. Reward anticipation was uniquely disrupted in stress-susceptible but not stress-resilient mice, compared to non-defeated controls, but such deficits depended upon whether or not another reward was recently obtained as well as the level of economic demand of the task. These findings suggest that the ability to detect motivational deficits after chronic stress is state-specific and can be masked by environmental pressure.

## Results

Following exposure to chronic social defeat stress (10 consecutive days, Fig. 1a-b), C57BL/6J male mice were categorized into either stress-susceptible (SUS) or stress-resilient (RES) groups based on a brief social interaction assay that has served as a well-validated predictor of several depression-related phenotypes on other rapid screening tests and on some more extensive behavioral procedures (Fig. 1d) (*22–29*). Next, these animals alongside non-defeated controls (CON) were tested across 55 consecutive days on Restaurant Row longitudinally (Fig. 1a), during which animals spend a limited time budget foraging for their only source of food (Fig. 1c) (*12–15*). On this task, mice make serial decisions accepting or rejecting offers presented for food rewards. The maze is divided into four uniquely flavored and contextualized feeding sites, or “restaurant” patches. Flavors are used to manipulate reward value without assuming individual differences in subjective preferences and without introducing variable handling time (e.g., as opposed to introducing different reward sizes). Each restaurant contains a separate offer zone and wait zone. Upon entry into a restaurant’s offer zone, a tone sounded in the offer zone whose pitch indicated the delay mice would have to wait in a cued countdown in order to earn a pellet should mice choose to enter the wait zone. After consuming an earned reward in the wait zone or skipping an offer in the offer zone, mice were required to advance to the next restaurant.

**Figure 1.**
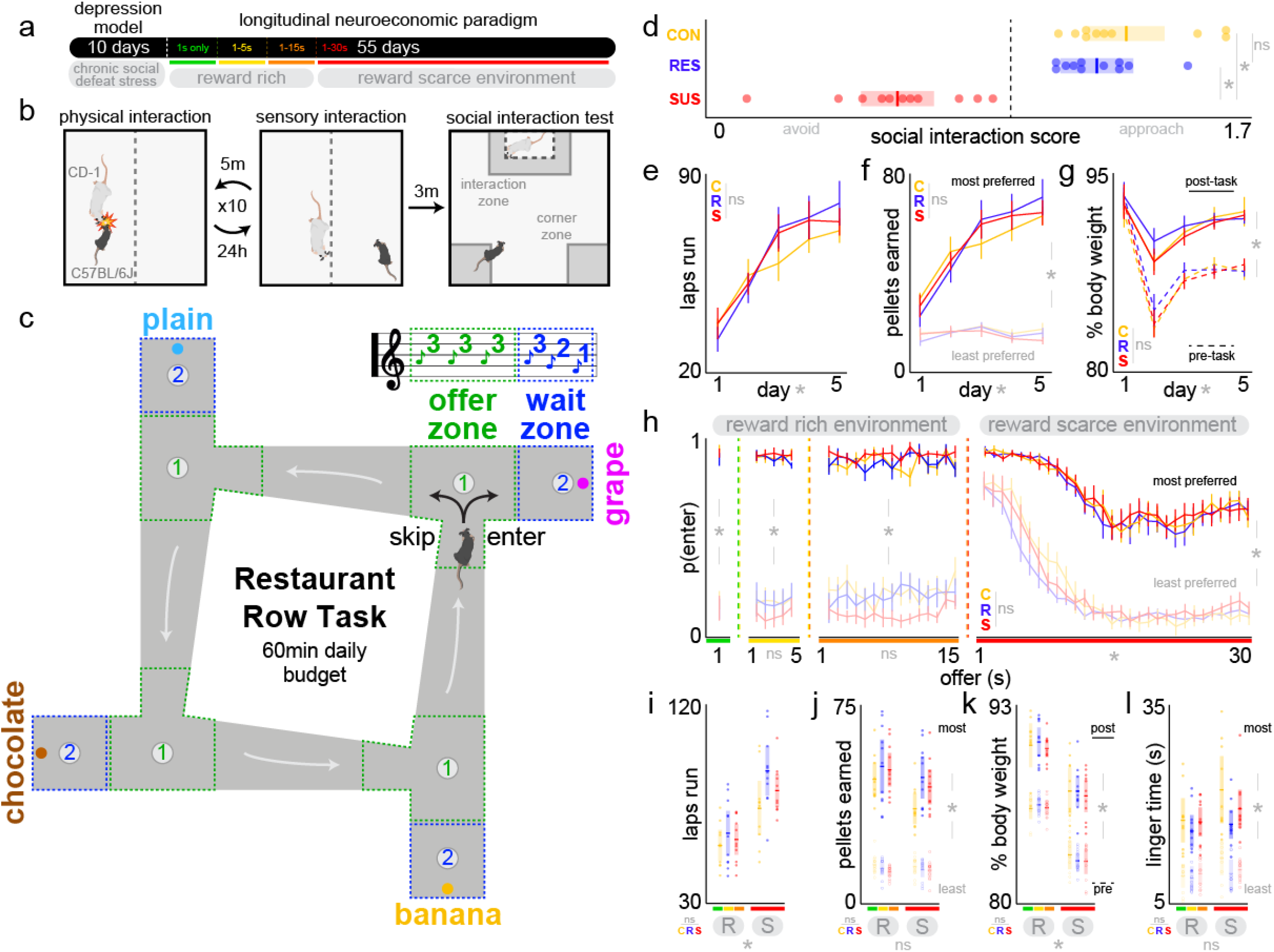
Mice exposed to chronic social defeat stress are capable of learning the Restaurant Row task without overt differences in revealed subjective flavor preferences, economic decisions, or post-consumption hedonic valuations. (a) Experimental timeline. (b) Chronic social defeat stress protocol. C57BL/6J mice are exposed to different aggressive CD-1 mice for 10 consecutive days (5-minute daily physical interactions followed by co-housing with a divider for the rest of the day). After the final defeat exposure, C57BL/6J mice are screened on the social interaction test. Time spent in the interaction zone is quantified before (3-minute baseline probe) and after (3-minute test probe) a novel CD-1 target mouse is placed in a small interaction chamber. (c) Restaurant Row task schematic. Mice have a limited time budget traversing a square maze with four uniquely flavored and contextualized restaurants, making serial accept versus reject decisions encountering offers randomly selected from a uniform distribution of delayed rewards cued by tone pitch in the offer zone that counts down should mice choose to enter the wait zone. (d) Social interaction score derived during the social interaction test is defined by ratio of time spent in the interaction zone when the CD-1 target is present versus when absent. Vertical dashed line represents score of 1.0. Data below this line reflect avoidance behavior that was used to define stress-susceptible (SUS, n=10) mice versus stress-resilient (RES, n=11) mice who instead approach the novel CD-1 target mouse during the social interaction screen, like non-defeated controls (CON, n=11). One-way ANOVA: *F*_2,31_=47.975, \**p*<0.0001, Tukey_CON/RES_ *t*=1.15, ^ns^*p*=0.495. (e) Laps run in the correct direction. Two-way ANOVA: Significant main effect across days: *F*_4,31_=81.953, \**p*<0.0001; no significant interaction with group: *F*_8,159_=1.676, ^ns^*p*=0.19. (f) Pellets earned in most preferred and least preferred restaurants. Rankings defined by summing end of session earns. Three-way ANOVA: Significant interaction between day and rank: *F*_4,139_=78.771, \**p*<0.0001; no significant interaction between day, rank, and group: *F*_4,319_=1.207, ^ns^*p*=0.30. (g) Percent bodyweight measured before and after testing relative to baseline weight defined before day 1 of Restaurant Row testing. Three-way ANOVA: Significant interaction between day and pre-vs. post-task measurements: *F F*_4,319_=9.966, \**p*<0.01; no significant interaction between day, pre-vs. post-task measurements, and group: *F*_4,319_=0.017, ^ns^*p*=0.98. (h) Probability of accepting an offer as a function of cued offer cost in a reward rich environment (1 s only [green], 1 to 5 s [yellow], and 1 to 15 s [orange] offers, note mice accept every offer in this environment in the most preferred restaurant regardless of cost) versus a reward scarce environment (1 to 30 s [red] offers). Four-way ANOVA: Significant interaction between offer, rank, and block of training: *F*_87,7038_=6.476, \**p*<0.01; no significant interaction between offer, rank, block of training, and group: *F*_174,7038_=0.001, ^ns^*p*=0.99. (i-l) Behaviors compared between reward rich (R) and reward scarce (S) environments: (i) laps run in the correct direction. Two-way ANOVA: Significant main effect across environments: *F*_1,31_=42.850, \**p*<0.0001; no significant interaction with group: *F*_2,63_=0.910, ^ns^*p*=0.41. (j) Pellets earned in most and least preferred restaurants. Three-way ANOVA: No significant main effect across environments: *F*_1,31_=0.012, ^ns^*p*=0.91; significant main effect of rank: *F*_1,31_=524.843, \**p*<0.0001; no significant interaction between environment, rank, and group: *F*_2,127_=0.456, ^ns^*p*=0.63. (k) Percent baseline bodyweight measured before and after testing. Three-way ANOVA: Significant main effect across environments: *F*_1,31_=188.431, \**p*<0.0001; significant main effect of pre-vs. post-task measurements: *F*_1,31_=312.3 10, \**p*<0.0001; no significant interaction between environment, pre-vs. post-task, and group: *F*_2,127_=0.077, ^ns^*p*=0.93. (l) Time spent lingering at the reward site after consuming an earned reward before advancing to the next trial in most and least preferred restaurants. Three-way ANOVA: No significant main effect across environments: *F*_1,31_=2.034, ^ns^*p*=0.50; significant main effect of rank: *F*_1.31_=83.936, \**p*<0.0001; no significant interaction between environments, rank, and group: *F*_2.127_=0.118, ^ns^*p*=0.89. Error bars represent ±1 SEM. Dots represent individual animals.

On days 1-5 of the task, all offers were only 1 s in length. During this time, mice displayed stable subjective flavor preferences that were ranked by summing end of session earnings in each restaurant and were no different on average between CON, RES, and SUS mice (Fig. 1f), running a similar number of laps (Fig. 1e) and stabilizing at equivalent body weights (Fig. 1g). Next, mice advanced into an increasingly reward scarce environment every five days, being exposed to a more expensive range of offers (days 6-10: 1 to 5 s offers; days 11-15: 1 to 15 s offers, days 16+: 1 to 30 s offers, Fig. 1a). While the probability of entering the wait zone was consistently higher for more preferred restaurants compared to less preferred restaurants across the entire experiment, mice did not factor in the cued cost of the offer in their offer zone decisions until entering the 1 to 30 s block of testing (Fig. 1h). CON, RES, and SUS mice did not differ in their ability to effectively discriminate tones in the offer zone during this block. By adopting this change in strategy in a reward scarce environment, mice were able to maintain food intake (Fig. 1j) but in turn expended more energy by running more laps (Fig. 1i) and thus had lower body weights (Fig. 1k), with no differences between CON, RES, and SUS mice. Furthermore, by measuring the time after consuming earned rewards until departing the wait zone, we found that mice lingered at the reward site before advancing to the next trial, lingering longer in more preferred restaurants (Fig. 1l). This metric represents a post-consumption within-trial place-preference associated with the restaurant’s context that was stable across the entire experiment and was not different among CON, RES, and SUS mice.

While there were no overt differences among CON, RES, and SUS mice in several measures of reward value, how mice approached each restaurant revealed other aspects of motivation that were different between groups. Inter-trial travel time was measured in the corridor between restaurants either from when a skip decision was made in the offer zone or from when mice departed the wait zone after consuming an earned reward. By analyzing travel time following skip decisions in a reward rich environment, we found that all mice arrived at the next restaurant more quickly when departing the least preferred restaurant compared to the most preferred restaurant (Fig. 2a). These data suggest that travel time captures a form of reward anticipation and invigorated motivational state in between trials based solely on the identity of the upcoming restaurant signaled by spatial context and independent of the delay of the next reward offer, which is not cued until arriving at the next restaurant.

**Figure 2.**
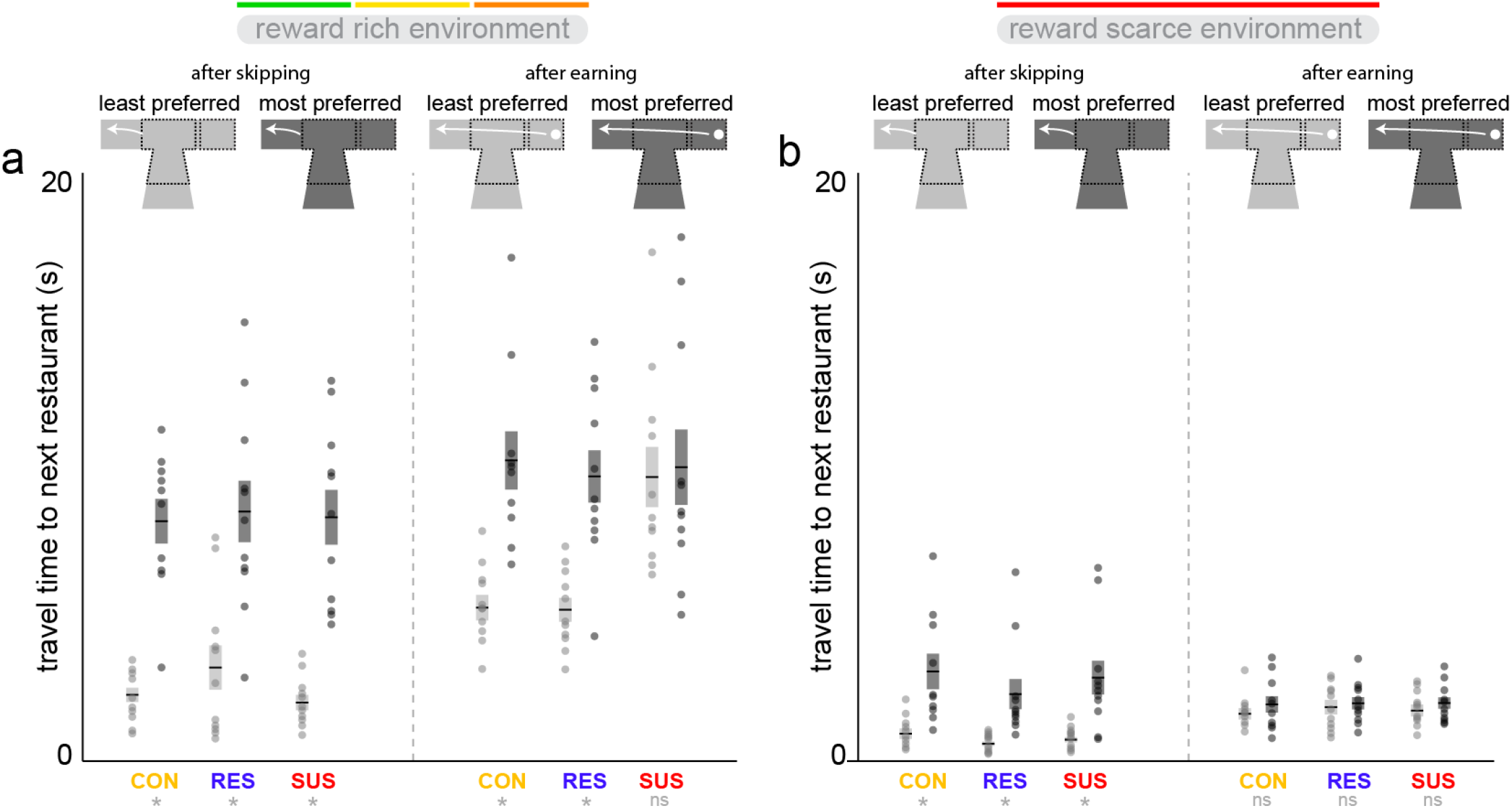
Travel time between restaurants reflects anticipatory behavior with a deficit in SUS mice that is state-specific and sensitive to economic pressure. (a) Travel time between restaurants in a reward rich environment (1 s only [green], 1 to 5 s [yellow], and 1 to 15 s [orange] offers) after a skip decision was made in the offer zone (left) versus after an earned reward was consumed and mice departed the wait zone (right) separated by the ranking of which restaurant is being departed. Overall, mice travel faster to the next restaurant after departing the least preferred restaurant (light gray) compared to after departing the most preferred restaurant (dark gray), main effect of rank: *F*_1,31_=90.264, \**p*<0.0001. Despite all animals showing this invigorated behavioral response after skipping a previous offer in the least preferred restaurant compared to the most preferred restaurant, only SUS mice do not display this invigorated behavioral response after earning rewards (significant interaction between rank, enter vs. skip decision on the previous trial, and group: *F*_2,127_=3.318, \**p*<0.05). (b) Travel time between restaurants in a reward scarce environment (1 to 30 s [red] offers). Overall mice travel faster in a reward scarce environment compared to a reward rich environment: *F*_1,31_=87.189, \**p*<0.0001. In this environment, invigorated travel times were observable after skipping the least preferred restaurant compared to after skipping the most preferred restaurant (left, *F*_1,31_=35.811, \**p*<0.0001), with no interaction with group (*F*_2,63_=0.202, ^ns^*p*=0.82). However, in this environment, there were no detectable differences in travel times observed after earning in either the least or most preferred restaurants in all groups (right, no main effect of rank: *F*_1,31_=2.060, ^ns^*p*=0.16, no interaction with group: *F*_2,63_=0.135, ^ns^*p*=0.87).

Next, by analyzing travel time after earns, overall travel times were slower compared to after skipping. Nonetheless, anticipatory effects were still measurable, with faster travel times once departing after earning in least preferred restaurants compared to most preferred restaurants in CON and RES mice (Fig. 2a). Interestingly, this effect was absent in SUS mice such that travel times were just as slow when departing least preferred restaurants after consuming an earned reward compared to when departing most preferred restaurants. These data indicate that anticipatory responses of more preferred future reward opportunities in SUS mice is blunted, not globally because anticipation is intact when skipping, but only upon reward receipt in least preferred restaurants. This suggests that impairments in eagerness to travel in SUS mice is state-specific. “State” here is defined by whether or not food was consumed on the previous trial. This finding highlights how the receipt of a reward only in SUS mice serves as an experience capable of devaluing the anticipation of better, future rewards while other animals under similar circumstances are able to invigorate their behavioral response in pursuit of upcoming opportunities.

Compared to behavior in a reward rich environment, travel time in a reward scarce environment overall was significantly faster (Fig. 2b) as mice were pressured to run more laps and make faster, more economically advantageous choices similar for CON, RES, and SUS mice. Despite this overall increase in speed, anticipation was still observable when skipping least preferred restaurants compared to most preferred restaurants in CON, RES, and SUS mice. Interestingly, anticipation after consuming an earned reward was no longer detectable in any group. These data suggest that the level of external economic pressure placed on animals can increase demand on foraging locomotion. This pressure can in turn mask state-specific anticipation and conceal potential stress-induced deficits in motivation.

## Discussion

Anticipating a reward can alter behavior in numerous ways, from increasing salivation in classical Pavlovian conditioning to invigorating more complex goal-oriented actions through instrumental transfer (*30, 31*). Deficits in reward anticipation can lead to impairments in motivation often observed in stress-related disorders such as depression, but it is unclear if such changes are linked to the intrinsic value of the reward itself, work required to obtain the reward, or other processes reflecting interactions between various internal states and external circumstances (*32*). We measured reward anticipation by examining travel behavior between serial foraging decisions, which is often formalized into neuroeconomic models of energetic optimization but is typically held constant, controlled for, or ignored in lab experiments rather than viewed as a dependent measure of interest (*33–37*). Here, we report stress-induced deficits in eagerness to travel that can be isolated in space and time from other decision-or consummatory-related valuations measured within the same trial and are dependent on reward history and environmental pressure.

Invigorated locomotion has previously been used to capture motivation but is often measured in tasks that do not parametrically vary value along multiple dimensions or different action-selection processes (*2, 38*). Furthermore, such tasks typically measure the pursuit of a single reward in an operant box with anticipation behavior operationalized by the degree of lever pressing, nose poking, or lick rate at a reward port (*39–43*). These metrics are inherently linked to preparatory consummatory behavior, goal tracking, or temporal error of estimated reward delivery (*44, 45*). Nonetheless, investigating the neural correlates of such processes have revealed a critical role of dopaminergic transmission interacting with reward circuitry and psychomotor systems. Dopamine signaling has been shown to respond to a variety of events on a spectrum from pure reward to pure movement, including unconditioned reward consumption, cue-related reward prediction, movement-locked reward-seeking behavior, generalized reward-related invigoration, and strict locomotor movements, broadly tuned to a variety of actions and typically bursting just before acceleration (*46–54*). Dopamine signaling is thought to gate the excitability of striatal output neurons that integrate cortical and thalamic reward-related action plans from coincident glutamatergic inputs translated into motor events (*55, 56*). Although much of the literature on dysfunction in such processes has traditionally focused on pathology involved in Parkinson’s disease, schizophrenia, and addiction, several studies have explored how alterations in dopamine transmission may underlie motivational deficits in stress-related disorders (*57*). Blunted reward prediction error signals in the striatum and ventral tegmental area have been observed in patients with major depressive disorder as well as in animals following exposure to stress (*26, 58–61*). Acute stress has been shown to alter the neural encoding of reward anticipation associated with diminished anticipatory licking behavior (*39*). Chronic social defeat stress alters dopaminergic firing rates in SUS mice, linked to anhedonia-related phenotypes but not reward anticipation processes specifically (*26, 60*). In the present study, why anticipatory behavior in SUS mice remains intact when skipping but not after recently earning a reward is perplexing. We suggest that state-specific differences here likely involve other neurotransmitter systems such as serotonin, which can signal quiescence of reward-related vigor following food consumption and have more classically been implicated in depression (*62–67*). Consuming a reward may involve a “reset” in relative value state from which a stress-related deficit emerges that is not present in the computations during skipping (Fig. 3). Dopamine and serotonin transmission is likely interacting here, both of which have been shown to contribute to more complex foraging strategies that incorporate travel speed, hunger, energy optimization, and opportunity costs when exploring an environment (*68, 69*). Given enough task pressure, decision history may have negligible effects on the physiological processes underlying movement vigor and may be less sensitive to reveal stress-related dysfunction.

**Figure 3.**
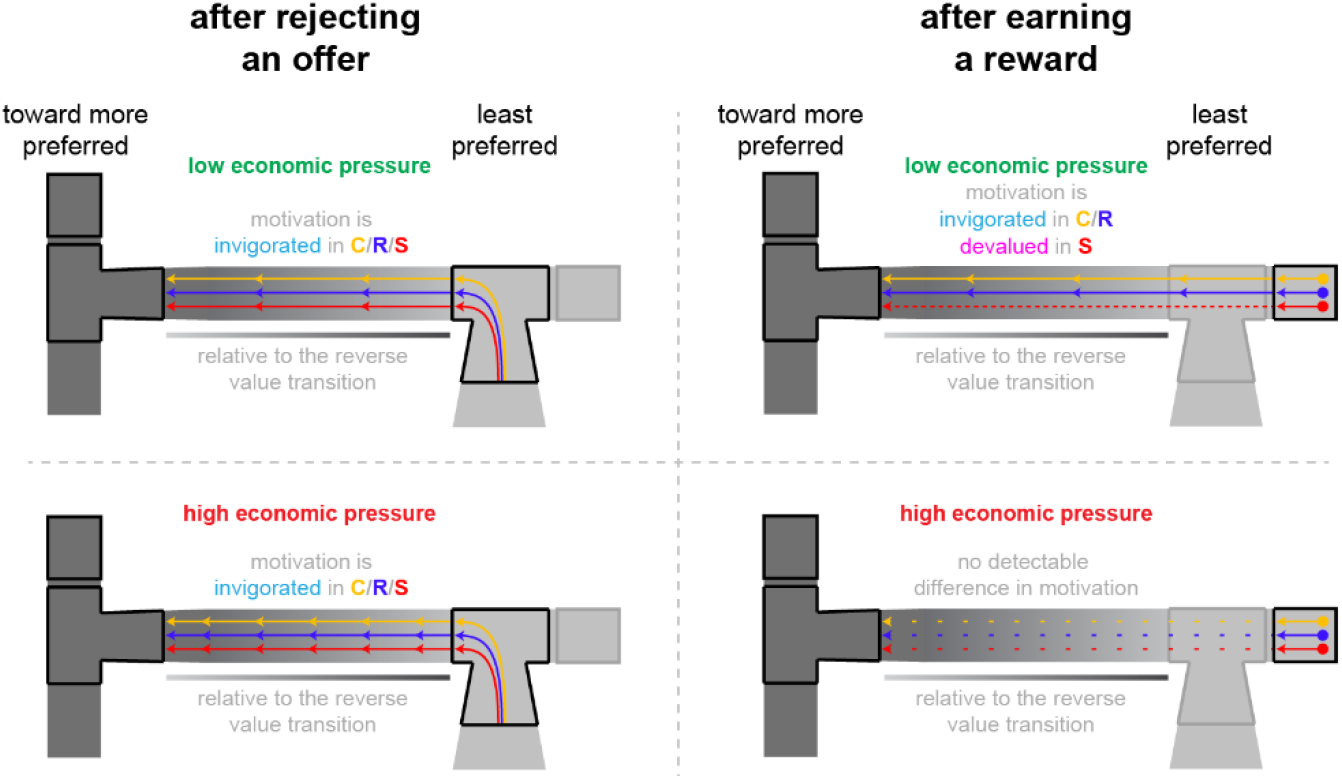
Summary of depression-related deficits in reward anticipation that are state-specific and masked by economic demand. Travel time between restaurants when traveling from the least preferred restaurant to more preferred restaurants, defined by rankings of total earns in each restaurant, and depicted by the transition in contrast from light to dark gray, is decreased (or invigorated, represented by the chevrons) relative to departures in the reverse value sequence (when traveling from the most preferred restaurant to less preferred restaurants, represented by the thin gradient bar from dark to light gray located beneath the corridor). This anticipatory effect is present in non-defeated control mice (C), stress-resilient mice (R), and stress-susceptible mice (S) after skipping an offer presented in the least preferred restaurant while in an economically rich environment (low economic pressure, top left). This anticipatory effect is also present in C and R mice after earning a reward in this environment (chevrons present), but is absent in S mice (chevrons absent, top right). In an economically scarce environment (high economic pressure, bottom), all mice travel faster (represented by the increase in chevrons). Anticipatory effects are still present in C, R, and S mice after skipping the least preferred restaurant (bottom left) but are no longer detectable in any group (represented by the dashed lines, increased spacing reflects the overall faster travel speed in this environment, bottom right).

We isolated a measure of reward anticipation based on travel vigor during a neuroeconomic task. We discovered that stress-susceptible but not stress-resilient mice show impairments in this measure of reward anticipation not merely as a result of a global deficit in anticipation capabilities but rather only following reward consumption. These data provide a novel perspective to interrogate how a rewarding stimulus only in a depressive state could devalue the anticipation of future rewards (*70, 71*). We also highlight the importance of careful considerations of task demands in being able to extract stress-related differences in motivation. Our approach can shed light on more nuanced aspects of anhedonia, unpacking how state and environment interact in certain circumstances to influence the way depressed individuals struggle with motivational inertia and perceive what may lie on the horizon ahead.

## Acknowledgments

We thank members of the Nestler and Russo labs for helpful discussion and technical assistance. Funding provided by: P50MH096890 (EJN), R01MH051399 (EJN), R01MH114882 (SJR), R01MH127820 (SJR), R01MH104559 (SJR). Open-source illustrations obtained from SciDraw (www.scidraw.io), credit Federico Claudi. We also thank Alexxai Kravitz for assistance in developing the open-source pellet dispensers used in this experiment (hackaday.io/project/171116-fed0).

## Author contributions

Conceptualization: BMS; Methodology: RDC, SJR, EJN, BMS; Investigation: RDC, FMR, LL, AMT, BMS; Data curation: RDC, BMS; Formal analysis: RDC, BMS; Funding acquisition: SJR, EJN; Supervision: RDC, SJR, EJN, BMS; Writing – original draft: RDC, BMS; Writing – review & editing: all authors

## Data and materials availability

All data, code, and materials used in the analysis are available in the materials and methods section or upon request.

## Materials and methods

### Animals and husbandry

We purchased 10-week-old wild-type male C57BL/6J mice (Jackson Laboratory). Additionally, we purchased 16 to 24-week-old male sexually experienced retired breeder CD-1 (ICR) mice (Charles River Laboratories) that were used as aggressors for the chronic social defeat stress protocol. All mice were maintained on a 12-hr light/dark cycle with ad libitum access to water. Experiments were conducted during the light phase. Experiments were approved by the Mount Sinai Institutional Animal Care and Use Committee (IACUC; protocol number LA12-00051) and adhered to the National Institutes of Health (NIH) guidelines.

### Chronic Social Defeat Stress

A single C57BL/6J mouse was co-housed with a single CD-1 mouse and allowed to experience aggression behavior for 5-10 min of attacking before being separated by a mesh divider for the remainder of the day. Mice then no longer had direct physical contact but continued to have visual, olfactory, and auditory contact. This was repeated for 10 consecutive days with 10 different CD-1 mice. Non-defeated control C57BL/6J mice were exposed instead to other, domiciled C57BL/6J mice. Mice had access to regular chow ad libitum during this protocol and were weighed daily.

### Social Interaction Screen

After the defeat protocol, mice were individually housed and assayed on a social interaction test that can capture social avoidance. A single C57BL/6J mouse was placed in a large open field arena with a novel CD-1 mouse enclosed in a small chamber. EthoVision software was used to track the location of the C57BL/6J mouse during this social interaction assay. Time spent near (interaction zone) versus away from the CD-1 mouse was used to quantify a social interaction score calculated from time in the interaction zone with the CD-1 mouse present in the chamber (2.5-min trial) relative to a preceding 2.5-min baseline trial without the target CD-1 mouse present.

### Neuroeconomic Decision-Making Paradigm: Restaurant Row Task

After the social interaction screen, mice were switched to a full-nutrition flavored pellet diet (BioServe products; 20 mg dustless precision full-nutrition pellets; a ~3 g mixture of chocolate, banana, grape, and plain flavored pellets as a daily ration) and food restricted to approach 80-85% of their free-feeding body weight over the next 3 days before starting training on the neuroeconomic operant decision-making paradigm termed “Restaurant Row.” Mice were weighed twice daily for the remainder of the experiment, before and after each Restaurant Row testing session. Mice were tested on Restaurant Row 7 days a week since performance on this task served as their sole source of food in this closed economy system. Mice foraged daily in a square maze for food rewards of varying cost (delays cued by tone pitch) and flavor (signaled by spatial context) while on a daily limited time budget (60 min). Each uniquely flavored “restaurant” was spatially fixed in the maze with patterns on the wall to signify the restaurant identity (chocolate: vertical stripes; banana: checkers; grape: triangles; plain, horizontal stripes). Rewards were delivered using a 3D printed automated pellet dispenser that was triggered by a computer running the behavioral task programmed in the ANY-Maze software made by the Stoelting Company. Behavioral events were triggered by spatial movements through the maze video-tracked by ANY-Maze. ELP USB camera with a Xenocam 1/2.7” 3.6 mm lens was used for video tracking. Restaurant Row testing took place in dim lighting conditions. The receptacle of the pellet dispenser also featured a custom-built trap door that would discard an uneaten pellet triggered upon exit from the wait zone if mice did not immediately consume food off of the pedestal. This prevented mice from hoarding rewards and forced animals to adhere to the structure of the task to make meaningful and intentional foraging decisions. Small wall mounted speakers (MakerHawk 3 Watt 8 Ohm Single Cavity Mini Speakers driven by a DROK 5W+5W Mini Amplifier Board PAM8406 DC 5V Dual Channel Class D) were fixed to the wall of each restaurant that played a 500 ms tone upon entry into the offer zone and repeated every s until either an enter or skip decision was made. The pitch of the tone varied depending on the randomly selected offer of that trial (1 s = 4,000 Hz and each second above that was an additional 387 Hz; e.g., 5 s = 5,548 Hz; 15 s = 9,418 Hz; 30 s = 15,223 Hz). Upon entry into the wait zone, the tones descended in a countdown fashion stepping down 387 Hz each second until a reward was earned and delivered. There is no penalty to quitting during the countdown other than the offer was rescinded and the mouse must advance to the next restaurant.

